# Comparative Analysis of Gene Expression Identifies Distinct Molecular Signatures of Bone Marrow- and Periosteal-Skeletal Stem/Progenitor Cells

**DOI:** 10.1101/210435

**Authors:** Lorenzo Deveza, Laura Ortinau, Kevin Lei, Dongsu Park

## Abstract

Periosteum and bone marrow (BM) both contain skeletal stem/progenitor cells (SSCs) that participate in fracture repair. However, the functional difference and selective regulatory mechanisms of SSCs in different location are unknown due to the lack of specific markers. Here, we report a comprehensive gene expression analysis of bone marrow SSCs (BM-SSCs), periosteal SSCs (P-SSCs), and more differentiated osteoprogenitors by using reporter mice expressing Interferon-inducible *Mx1* and *Nestin*^GFP^, previously known SSC markers. We first defined that the BM-SSCs can be enriched by the combination of *Mx1* and *Nestin*^GFP^ expression, while endogenous P-SSCs can be isolated by positive selection of *Mx1*, CD105 and CD140a (known SSC markers) combined with the negative selection of CD45, CD31, and *osteocalcin*^GFP^ (a mature osteolineage marker). Comparative gene experession analysis with FACS-sorted BM-SSCs, P-SSCs, *Osterix*^+^ (OSX) preosteoblasts, CD51^+^ stroma cells and CD45^+^ hematopoietic cells as controls revealed that BM-SSCs and P-SSCs have high similarity with few potential differences without statistical significance. We also found that CD51^+^ cells are highly heterogeneous and little overlap with SSCs. This was further supported by the microarray cluster analysis, and the two populations clustered together. However, when comparing SSC population to controls, we found several genes that were uniquely upregulated in endogenous SSCs. Amongst these genes, we found KDR (aka Flk1 or VEGFR2) to be most interesting and discovered that it is highly and selectively expressed in P-SSCs. This finding suggests that endogenous P-SSCs are functionally very similar to BM-SSCs with undetectable significant differences in gene expression but there are distinct molecular signatures in P-SSCs, which can be useful to specify P-SSC subset *in vivo*.

## Introduction

Bone fractures constitute a significant burden to the healthcare system with about 16 M fractures per year in the United States. Majority of fractures heal with adequate treatment, but about 5-10% go on to non-union [1]. Treatment methods include bone grafting, delivery of growth factors, and cell-based therapies [1,2]. Fundamentally, these attempts to augment the healing process are attempts to stimulate the cells that drive fracture repair. Studies on such therapeutic attempts are based on using or stimulating bone marrow skeletal stem/progenitor cells (BM-SSCs), also known as bone marrow mesenchymal cells (BMSCs) [3]. However, endogenous SSCs are heterogeneous population and are present in multiple tissue location including periosteum [4]. Despite SSCs are necessary for fracture repair, yet whether SSCs in different location have same functional properties or they have distinct function and regulation that are necessary of the repair process remain unknown.

At its core, bone fracture healing is a complex process that involves the interplay of multiple cell types derived from different tissue sources. Bone marrow (BM) and periosteum are two of the surrounding tissue intimately involved in fracture repair [5]. However, BM is not necessary for healing to proceed, while removal of periosteal tissues can cause non-union. Indeed, this is a fundamental principle in clinical fracture management [6]. This is further exemplified by cell-labeling studies demonstrating that the major cellular contribution to the fracture callus are periosteal cells [7]. More importantly, it has been reported that P-SSCs may have differing functions than BM-SSCs [6,8], whereby P-SSCs display endochondral ossification and intramembranous bone formation, while BM-SSCs only participate in the latter process [8]. These differences suggest that P-SSCs may have different inherent properties compared to BM-SSCs.

Although there has been extensive studies to define unique gene expression patterns in postnatal skeletal stem cells [9], to date, there have been no studies looking specifically into the potential differences between P-SSCs and BM-SSCs. This is partly because no reliable markers exist to isolate each of these cell populations to enable such study. Studies on mouse BM-SSCs have identified multiple markers that isolate a potentially more highly purified population of these cells, including *Nestin*^*GFP*^ [10], *LepR*^*Cre*^ (Leptin Receptor) [11], and *Grem1*^*Cre-ERT*^ (Gremlin 1) [12]. Previously, *Myxovirus resistance 1* (*Mx1*) was also shown to identify long-term resident skeletal stem/progenitor cells in mice via in vivo imaging experiments consistent with their role as BM-SSCs [13]. While fewer markers exist for P-SSCs, *Mx1*^+^ cells are known to reside within the periosteal compartment [13], and these cells also provide downstream osteolineage cells enabling their potential use for endogenous P-SSC study.

In this study, we isolate BM-SSCs and P-SSCs from transgenic mice based on expression of *Mx1* promoter. BM-SSCs were isolated from BM tissues in transgenic mice expressing *Mx1*^*Cre*^ and *Nestin*^*GFP*^(*Mx1*^+^*Nes*^GFP+^ cells), known SSC markers. P-SSCs were isolated from periosteal tissues in *Mx1*^Cre^; *ROSA*^*Tomato*^; *Osteocalcin*^GFP^ reporter mice, whereby P-SSCs were negatively selected against *Osteocalcin*^GFP+^ osteoblasts (*Mx1*^+^*Ocn*^−^ cells). Microarray was run on these cell populations, using CD45^+^ cells and Osterix (*Osx*^+^) osteolineage cells as controls. We further compared CD51^+^ cells as an additional BM-SSC population reported in literature. Lastly, we identify a potentially novel marker for mouse P-SSCs.

## Materials and Methods

### Mice

Four to six-week old C57BL/6, *Mx1*^*Cre*^ [14], *Rosa26-loxP-stop-loxP-tdTomato* (*Rosa*^*Tomato*^) mice were purchased from The Jackson laboratory. *Osteocalcin*^*GFP*^ [15] and *Nestin*^*GFP*^ [10] (C57/BL6 background) mice were kindly provided by Drs. Henry Kronenberg and Ivo Kalajzic. Genotyping of all Cre-transgenic mice and the Rosa locus was performed by PCR (GenDEPOT) according to The Jackson laboratory’s protocols. At 4-week age, all *Mx1* mice (*Mx1*^*Cre*^; *Rosa*^*Tomato*^; *Osteocalcin*^*GFP*^ *or Mx1*^*Cre*^; *Rosa*^*Tomato*^; *Nestin*^*GFP*^) were lethally irradiated with 9.5 Gy and transplanted with 10^6^ whole bone marrow cells from wild-type C57BL/6 mice (WT-BMT). At Six to eight weeks later (when host hematopoietic cells are less than 1%), *Mx1*^*Cre*^ activity was induced by intraperitoneal injection of 25 mg/kg of pIpC (Sigma) every other day for 10 days as described previously [10]. At the indicated time after pIpC induction, mice were subjected to in vivo imaging experiments. All mice were maintained in pathogen-free conditions, and all procedures were approved by Baylor College of Medicine’s Institutional Animal Care and Use Committee (IACUC).

### Intravital imaging

For *in vivo* imaging of fluorescent cells in living animals, mice were anesthetized with Combo-III and prepared for a customized two-photon and confocal hybrid microscope (Leica TCS SP8MP with DM6000CFS) specifically designed for live animal imaging, as described in our previous report [13,16]. Briefly, a small incision was introduced on the scalp of Mx1/Tomato/Ocn-GFP or Mx1/Tomato/Nestin-GFP mice and the surface of calvaria near the intersection of sagittal and coronal suture was exposed. The mice were then mounted on a 3-D axis motorized stage (Anaheim Automation Anaheim, CA), and the calvarial surface was scanned for second harmonic generation (SHG by femto-second titanium:sapphire laser pulses: 880 nm) from bones to identify the injury sites and the intersection of sagittal and coronal sutures. GFP-expressing cells (488 nm excitation, 505–550 nm detection) and Tomato-expressing cells (561 nm excitation, 590–620 nm detection) were simultaneously imaged by confocal spectral fluorescence detection. All images were recorded with their distances to the intersection of the sagittal and coronal sutures to define their precise location. After *in vivo* imaging, the scalp was closed using a VICRYL plus suture (Ethicon), and post-operative care was provided as previously described. 3-D Images were reconstructed using the Leica Application Suite software, and osteoblasts were counted.

### Isolation and flow cytometry analysis of mouse SSCs

To isolate periosteal cells, dissected femurs, tibias, pelvis and calvaria from mice were placed in PBS, and the overlying fascia, muscle, and tendon were carefully removed. The bones with periosteum were incubated in ice-cold PBS with 1% FBS for 30 min, and the loosely associated periosteum was peeled off using forceps, scalpel, and dissecting scissors. The soft floating periosteal tissues collected with a 40-µm strainer were then incubated with 5–10 ml of 0.1 % collagenase and 10% FBS in PBS at 37°C for 1 hour, and dissociated periosteal cells were washed with PBS, filtered with a 40-µm strainer and resuspended at ~1 × 10^7^ cells/ml. To isolate cells from bones and bone marrow, dissected femurs, tibias and pelvis bones after periosteum removal were cracked with a pestle and rinsed 3 times to remove and collect bone marrow cells. The remaining bones were minced with a scalpel and/or a dissecting scissor and then incubated with 10 ml of 0.1 % collagenase and 10% FBS in PBS at 37°C for 1 hour with strong vortexing every 10 minute. Dissociated cells were washed with PBS, filtered with a 40-µm strainer and resuspended at ~1 × 10^7^ cells/ml. To analyze or isolate SSCs and osteogenic cells, cells were stained with CD105-PE-Cy7 (clone: MJ7/18), CD140a-APC (clone: APA5), CD45-pacific blue (clone: 30-F11), Ter119-APC-Cy7 (clone: TER-119), and CD31-eFlour 450 (clone: 390) in combination with KDR-PE-Cy7 (clone: J073E5). Antibodies were purchased from eBioscience unless otherwise stated. Propidium iodide was used for viable cell gating. Flow cytometric experiments and sorting were performed using the LSRII and FACS Aria cytometer (BD Biosciences, San Jose, CA). Data were analyzed with the FlowJo software (TreeStar, Ashland, OR) and represented as histograms, contour, or dot plots of fluorescence intensity.

### Microarray analysis

Sorted cells pooled from five or more male and female mice were used to isolate RNA using the RNeasy Micro kit (Qiagen), according to the manufacturer’s instructions. Purified RNA was reverse-transcribed, amplified, and labeled with the Affymetrix Gene Chip whole transcript sense target labeling kit. Labeled cDNA (2 biological repeats) from indicated cells was analyzed using Affymetrix mouse A430 microarrays, according to the manufacturer’s instructions, performed at the Dana-Farber Cancer Institute Microarray Core. CEL files (containing raw expression measurements) were imported to Partek GS, and data were preprocessed and normalized using the RMA (Robust Multichip Average) algorithm.

### Microarray data analysis and statistics

Microarray data was pre-processed for normalization and statistical differences using R statistical package. Normalization was done using a robust multichip average (RMA) technique. Statistical differences were calculated with the limma package in R. Post-processing cluster analysis was done using Cluster 3.0 software and were plotted using Java TreeView software. Scatter plots were generated using Orange biolab software. We assessed pairwise comparisons between each of the following groups: 1) *Mx1*^+^Ocn^−^ P-SSCs; 2) *Mx1*^+^*Nes*^+^ BM-SSCs; 3) CD45^+^ hematopoietic lineage cells; 4) *Osterix*^GFP+^ osteoprogenitor cells [17]; and, 5) CD51^+^ BMSCs [18]. We evaluated the number of statistically different genes by changing the p-value statistical criteria for acceptance. We found that acceptance criteria of p < 0.05 provided at least 50 statistically different genes between controls and *Mx1*^+^ SSCs.

## Results

### *In vivo* dentification of BM-SSCs and P-SSCs

BM-SSCs and P-SSCs were derived from transgenic mice based on expression of interferon-induced GTP-Binding Protein *Mx1* promoter, which had been previously shown to represent long-term resident lineage restricted osteoprogenitor cells [13]. Here, *Mx1*^*Cre*^; *Rosa*^*Tomato*^; *Osteocalcin*^*GFP*^ mice were used as previously described [16]. Using this model, we confirmed through pulse-chase labeling studies in native bone marrow tissue (i.e. no injury) that pulse-labeled *Mx1*^+^ cells at day 5 are mainly *Osteocalcin*^GFP^ negative (*Ocn*^−^) and these *Mx1*^+^ cells contribute to the majority of new *Ocn*^+^ osteoblasts at day 60 (yellow), demonstrating that *Mx1*^+^ cells include skeletal stem/progenitor cells (SSCs), albeit far upstream of mature osteoblasts (Fig. 1A). Considering that *Ocn*^+^ cells represent mature osteolineage cells, we found that *Mx1*^+^Ocn^−^ upstream progenitors are present throughout bone marrow, as well as calvarial suture and periosteum (Fig. 1B). We thus isolated P-SSCs from periosteal tissue by focusing on *Mx1*^+^*Ocn*^−^ cells within this tissue compartment. Specifically, we isolated P-SSCs by isolating cells from periosteum, negatively selecting for hematopoietic lineage cells (CD45^−^), endothelial lineage cells (CD31^−^), erythroid lineage (Ter119^−^), and adult osteolineage cells (*Ocn*^−^), and positively selecting for SSC markers including *Mx1*^+^, CD105^+^ and CD140a^+^ (PDGFRa). We refer to these cells as *Mx1*^+^*Ocn*^−^ P-SSCs.

**Fig. 1.**
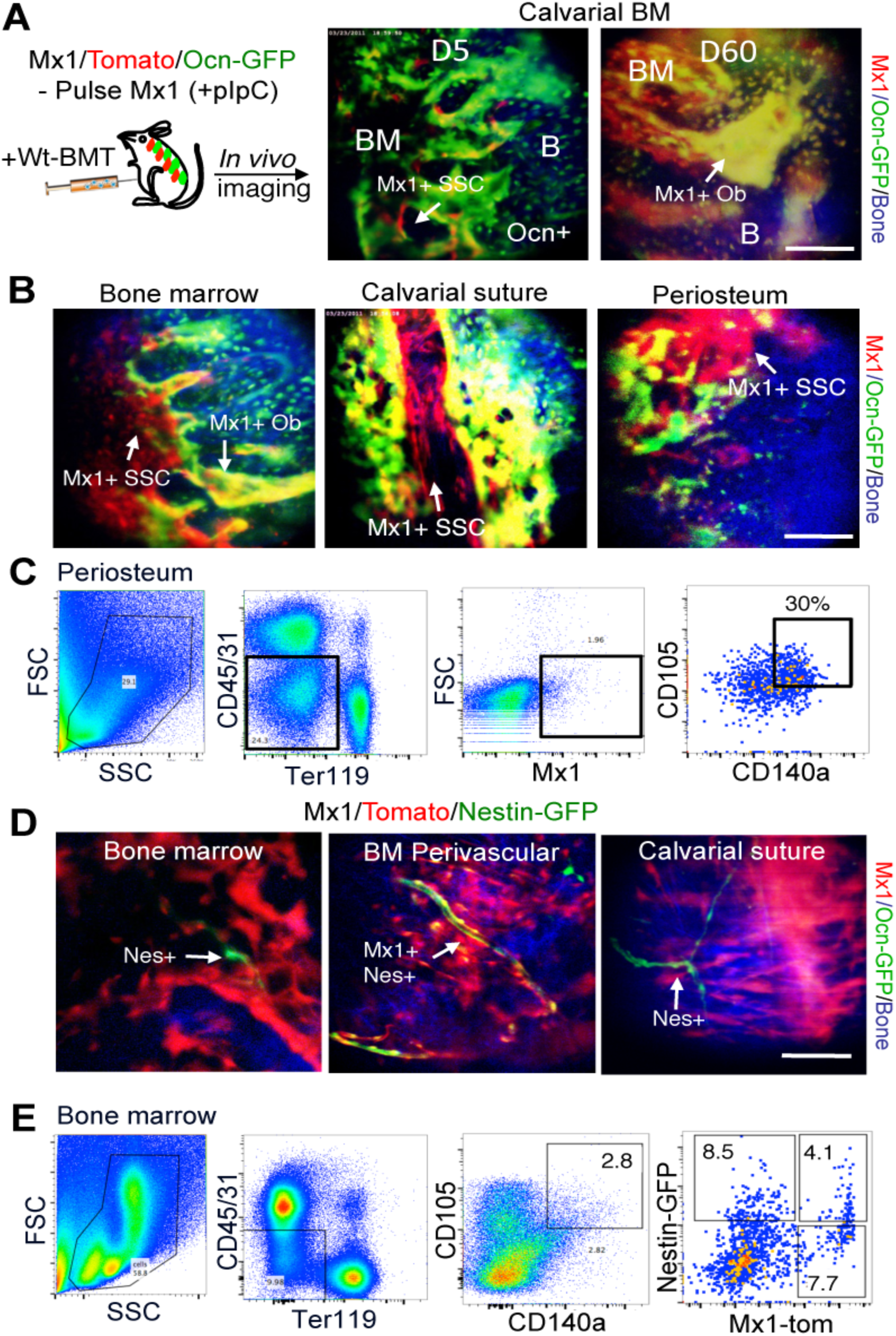
Functional identification of P-SSCs and BM-SSCs. (A.) Interferon inducible *Mx1*^+^ SSCs (red) are shown to contribute to majority osteoblasts (green, overlap yellow) *in vivo.* B) *Mx1*^+^ SSCs represent long-term osteolineage progenitor cells in BM and periosteal tissues. C) P-SSCs are derived from periosteal tissues and are FACS sorted by CD45^−^CD31^−^Ter119^−^ *Mx1*^+^*Ocn*^−^CD105^+^CD140a^+^, which are referred to as *Mx1*^+^*Ocn*^−^ P-SSCs. D) *Mx1*^+^*Nestin*^+^ BM-SSCs are perivascular cells in BM but are undetectable in periosteum and calvarial suture. E) Mx1^+^Nes^+^ cells within CD45^−^CD31^−^Ter119^−^CD105^+^CD140a^+^ SSC fraction in bone marrow are isolated by FACS-sorting and are referred to as *Mx1*^+^*Nes*^+^ BM-SSCs. Notably, CD105^+^CD140a^+^ progenitors are heterogeneous *Mx1*^+^ and *Nestin*^+^ cells.

BM-SSCs were isolated from *Mx1*^*Cre*^; *Rosa26*^*Tomato*^; *Nestin*^*GFP*^ transgenic mice. *Nestin*^GFP^ (*Nes*^+^) is a well-studied marker for BM-SSCs [10]. By pulse-chase labeling studies, we found that *Mx1*^+^*Nes*^+^ cells are native perivascular cells that are present throughout BM and calvarial suture (Fig. 1D), which is consistent with prior studies as BM-SSCs are generally known to be perivascular cells [10,19]. For our experiments, we isolate BM-SSCs using this model from the BM tissue compartment, which are sorted by negative selection of CD45, CD31, Ter119, as well as positive selection of CD105, CD140a (PDGFRa); finally, *Mx1*^+^*Nes*^+^ cells are selected from the remaining cells (Fig. 1E). We refer to this subpopulation as *Mx1*^+^*Nes*^+^ BM-SSCs. Interestingly, we noted that the selection of BM-SSCs based on CD45^−^CD31^−^Ter119^−^CD105^+^ CD140a^+^ cells yields a heterogeneous mixture of *Mx1*^+^ and *Nestin*^+^ cells (Fig. 1E).

### Common selection criteria for BM-SSCs yields a heterogeneous mixture

Microarray analysis was next performed on *Mx1*^+^*Ocn*^−^ P-SSCs and *Mx1*^+^*Nes*^+^ BM-SSCs to assess for functional genetic differences. We added an additional BM-SSC population that was selected from the BM compartment based on CD45^−^CD31^−^Ter119^−^CD105^+^CD140a^+^ selection, in addition to CD51^+^, which is a commonly used selection criteria for BMSCs [18]. We refer to these cells as CD51^+^ BMSCs. CD45^+^ hematopoietic lineage cells and *Osterix*^+^ (*Osx*^GFP+^) [17,20] osteoprogenitor cells were used as control populations. From scatter plot analysis of all microarrayed genes, we found that each SSC population is a distinct population as compared to CD45^+^ cells (Fig. 2A-C). We further found that each SSC population are similarly more closely related to *Osx*^+^ osteolineage cells, but with multiple differentially expressed genes (Fig. 2D-F). Taken together, these scatter plots illustrated that each SSC population is similarly distinct from CD45^+^ and *Osx*^+^ cells.

**Fig. 2.**
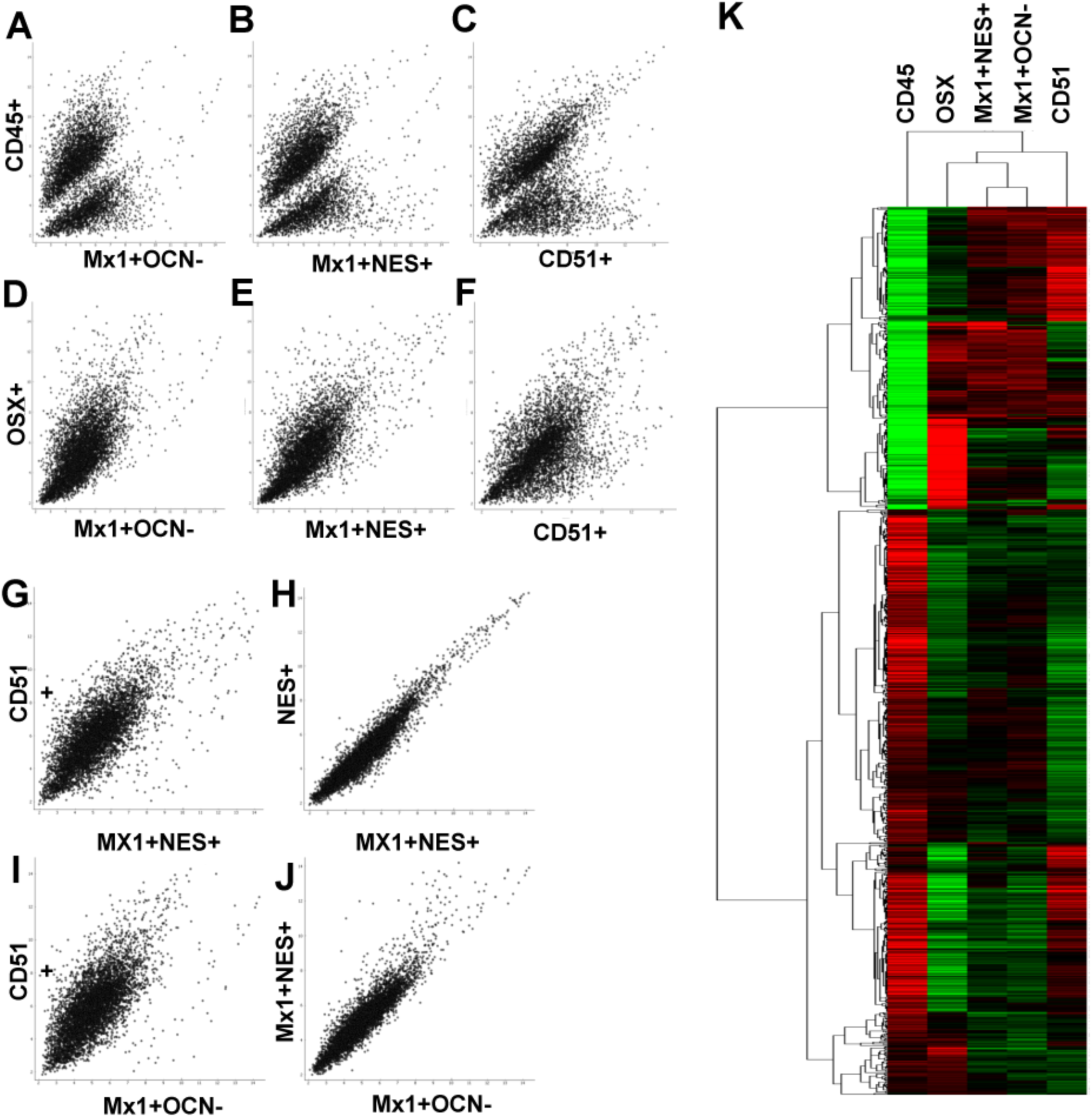
Commonly used markers for BM-SSCs yield a heterogeneous mixture, but are similar to P-SSCs. (A-C) Scatter plot comparison between *Mx1*^+^*Ocn*^−^ P-SSCs, (A), *Mx1*^+^*Nes*^+^ BM-SSCs (B), and CD51^+^ BMSCs (C) with CD45^+^ cells, demonstrates that these populations are likewise different from CD45^+^ cells within the BM compartment. (D-E) Scatter plot comparison between *Mx1*^+^*Ocn*^−^ P-SSCs, (D), *Mx1*^+^Nes^+^ BM-SSCs (E), and CD51^+^ BMSCs (F) with *Osx^+^ osteolineage cells shows that each of these populations are more functionally similar to the osteolineage cells. (G) Direct comparison between *CD51*^+^ BMSCs and *Mx1*^+^*Nes*^+^ BM-SSCs demonstrates that these two commonly used selection markers for BM-SSCs yield a heterogeneous mixture of cells. (H) *Mx1*^+^*Nes*^+^ BM-SSCs and *Nes*^+^ cells are essentially the same population of cells. (I-J) Comparing *Mx1*^+^Ocn*^−^ P-SSCs with CD51^+^ BMSCs (I) shows that these are functionally different cell-populations, but comparison with *Mx1*^+^*Nes*^+^ BM-SSCs (J) shows few differences. (K) Cluster analysis of these cell populations confirms scatter plot analysis and shows that *Mx1*^+^*Ocn*^−^ P-SSCs and *Mx1*^+^Nes^+^ BM-SSCs cluster together, but each of these populations are distinct from CD51^+^ BM-SSCs (p < 0.05).

Interestingly, we found that commonly used selection criteria for BMSCs may yield a heterogeneous mixture of cells, which is demonstrated by direct comparison between *Mx1*^+^Nes^+^ BM-SSCs and CD51^+^ BMSCs (Fig. 2G). Between these two cell populations there were 97 differentially expressed genes at p < 0.01 and 430 differentially expressed genes at p < 0.05. When comparing *Mx1*^+^*Nes*^+^ BM-SSCs with *Nes*^+^ BMSCs (i.e. *Mx1*^+/-^) there were no differentially expressed genes (Fig. 2H). These findings suggest that although both *Nes*^+^ and CD140a^+^CD51^+^ have both been shown to represent BMSCs, that BMSCs are a heterogeneous mixture of cells.

### P-SSCs and BM-SSCs are a similar population of cells

When directly comparing BM-SSCs with P-SSCs, we find that these cell populations are a similar population of cells. In our analysis, we found that CD51^+^ BMSCs had several differentially expressed genes compared to *Mx1*^+^*Ocn*^−^ P-SSCs (Fig. 2I), but there were few differences when comparing *Mx1*^+^*Nes*^+^ BM-SSCs with *Mx1*^+^*Ocn*^−^ P-SSCs and none were significant at the p < 0.05 acceptance criteria (Fig. 2J). This is further summarized in the cluster plot, which demonstrated that *Mx1*^+^*Nes*^+^ BM-SSCs and *Mx1*^+^*Ocn*^−^ P-SSCs clustered together and were separate from CD51^+^ BM-SSCs (p < 0.05, Fig. 2K).

### Determination of differentially expressed genes between P-SSCs and BM-SSCs with controls

Considering that there were no differentially expressed genes found between *Mx1*^+^Ocn^−^ P-SSCs and *Mx1*^+^*Nes*^+^ BM-SSCs, we proceeded to identify the genes that were differentially expressed between each of these populations and controls separately. Cluster analysis of differentially expressed genes between *Mx1*^+^*Ocn*^−^ and controls is shown in Fig. 3A and between *Mx1*^+^*Nes*^+^ BM-SSCs and controls is shown in Fig. 3B. There were 101 differentially expressed genes between *Mx1*^+^*Ocn*^−^ P-SSCs compared to controls and 84 for *Mx1*^+^*Nes*^+^ BM-SSCs; while there were 55 overlapping differentially expressed genes for both SSC populations compared to controls (Fig. 3C). Genes that were overexpressed are shown in Fig. 3D and Supplemental table 1. Amongst these genes, we were interested to find increased expression of the vascular endothelial growth factor receptors (VEGFR), Flt1 (VEGFR1) and KDR (VEGFR2), in the P-SSC population. Between these two genes, KDR was overexpressed in both SSC populations by the microarray analysis (Fig. 3D). The full list of differentially expressed genes is given in Supplemental table 1.

**Fig. 3.**
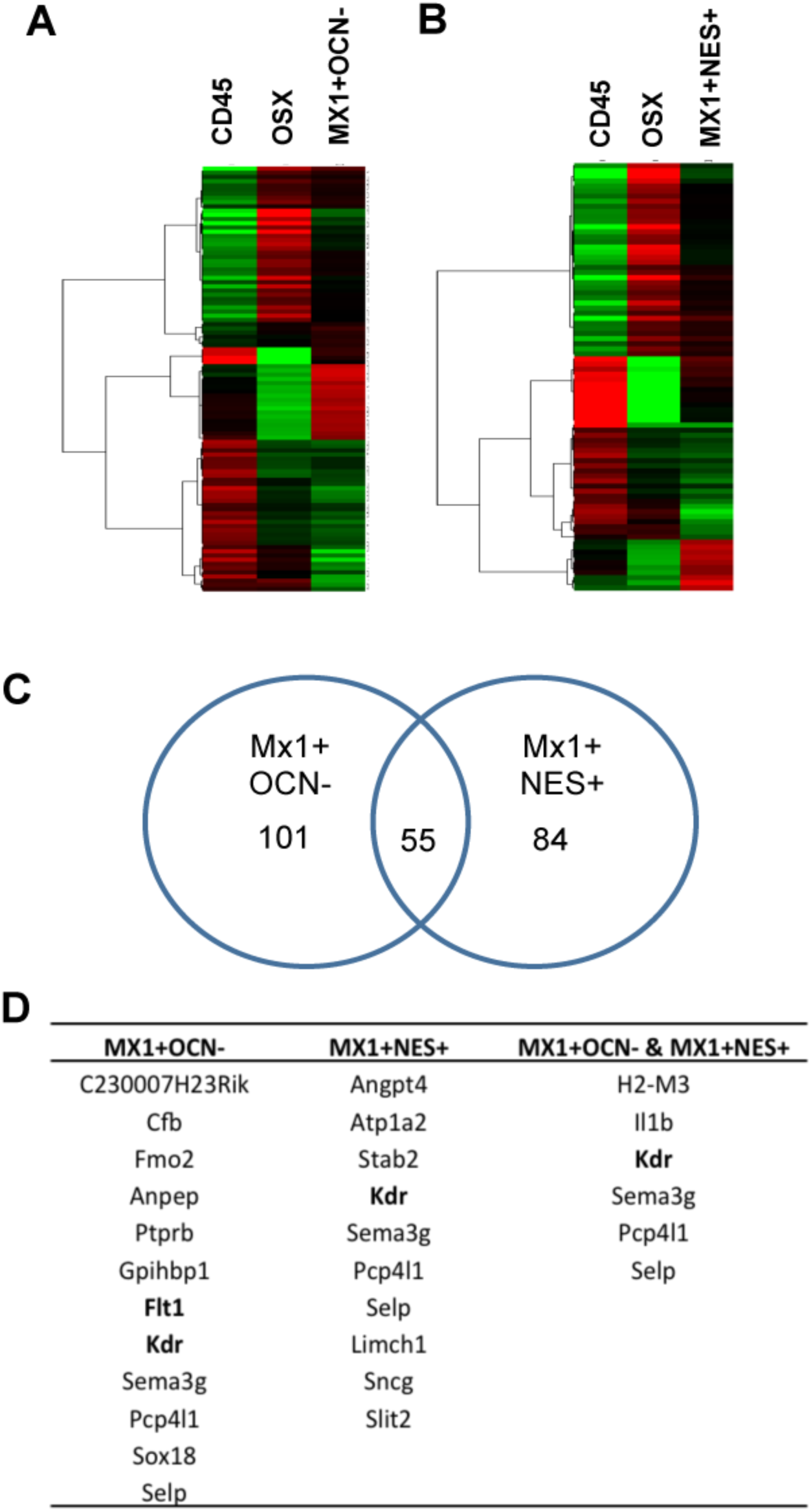
Identification of differentially expressed gene analysis between P-SSCs and BM-SSCs and controls. Differential gene expression between *CD45*^+^ cells and *Osx*^+^ cells with (A) *Mx1*^+^*Ocn*^−^ P-SSCs and (B) *Mx1*^+^*Nes*^+^ BM-SSCs. (C) Number of differentially expressed genes between SSC populations and controls shows 101 for *Mx1*^+^*Ocn*^−^ P-SSCs, 84 for *Mx1*^+^*Nes*^+^ BM-SSCs, and 55 overlap genes. (D) Table of genes that were upregulated in SSCs compared to controls shows some interesting vascular endothelial growth factor receptors (VEGF), including Flt1 (VEGF receptor 1) and KDR (VEGF receptor 2), despite removal of CD31 and Ter119 endothelial lineage cells from these populations.

### P-SSCs are CD140a^+^KDR^+^ stem/progenitor cells

From our microarray analysis, we sought to further explore KDR expression in *Mx1*^+^*Ocn*^−^ P-SSCs and *Mx1*^+^*Nes*^+^ BM-SSCs. Notably, CD140a^+^KDR^+^ cells have been found to represent early mesoderm subpopulations. We first compared our SSC populations to other publically available SSC populations using Gene Commons data (Fig. 4A). We noted that other well-studied BM-SSC markers, *Leptin receptor* (*Lepr*) and *Gremlin 1* (*Grem 1*), were highly expressed in *Mx1*^+^*Ocn*^−^ P-SSCs and *Mx1*^+^*Nes*^+^ BM-SSCs, which demonstrated the consistency of our data with other known SSC populations (Fig. 4A). By this same analysis, we found that KDR appeared to be higher expression in *Mx1*^+^*Ocn*^−^ P-SSCs than *Mx1*^+^*Nes*^+^ BM-SSCs, thereby supporting our microarray analysis. We next assessed KDR expression by FACS analysis (Fig. 4B-C). We included P-SSCs (CD45^−^CD31^−^Ter11^−^ CD105^+^CD140a^+^ *Mx1*^+^*Ocn*^−^), periosteal adult osteolineage cells (CD45^−^CD31^−^Ter119^−^*Mx1*^−^*Ocn*^+^), BMSCs (CD45^−^CD31^−^Ter119^−^CD140a^+^ *Nes*^+^), and CD45^+^ cells. We found that P-SSCs had increased expression of CD140a and KDR with 72% of the population overexpressing these markers and this was significantly increased compared to *Nes*^+^ BMSCs (n = 3, p < 0.0001, Fig. 4D). Thus, while our microarray analysis demonstrated increased KDR expression in both BM-SSCs and P-SSCs, we found via FACS analysis that P-SSCs have selectively high expression of KDR.

**Fig. 4.**
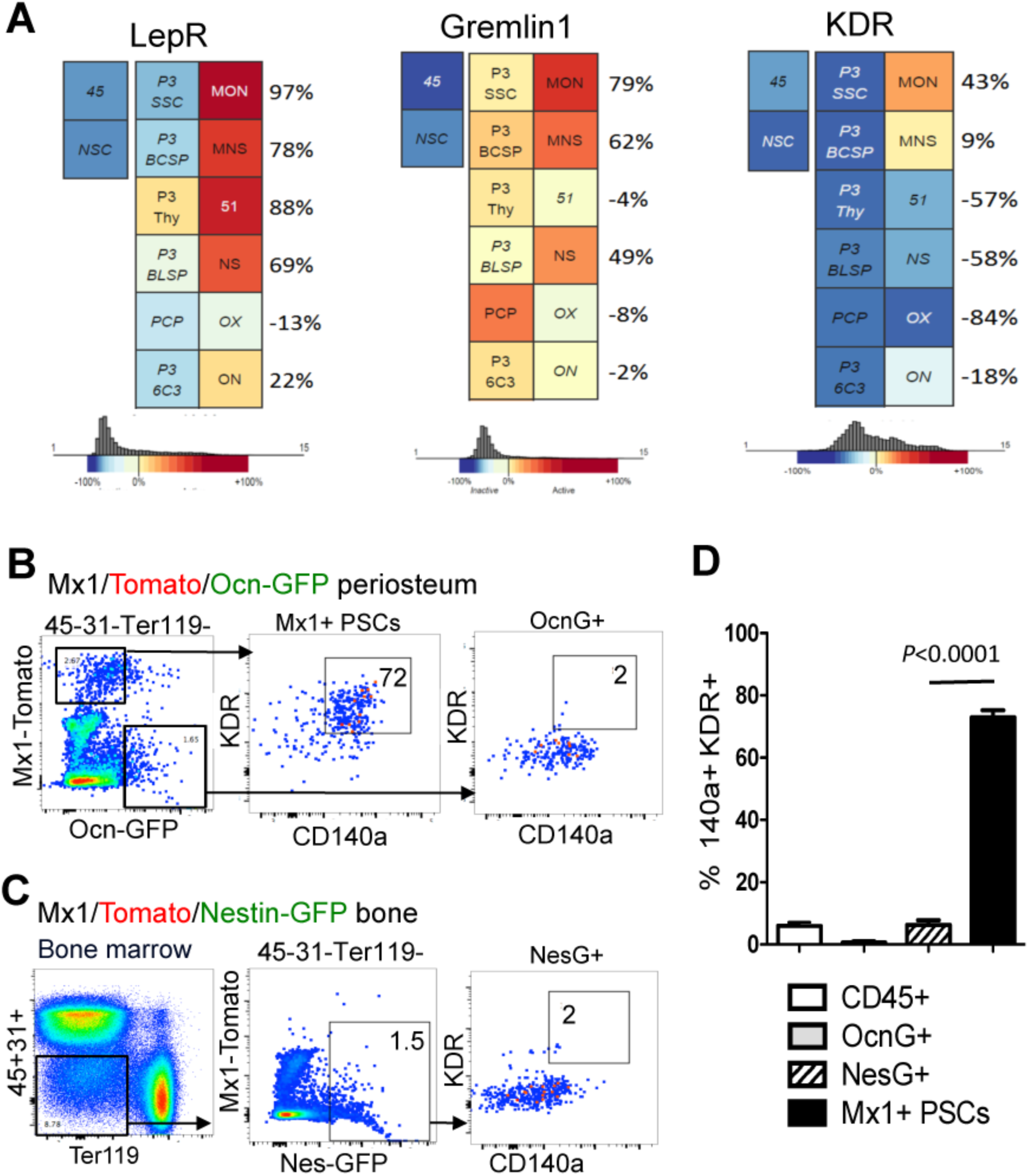
P-SSCs are KDR^+^CD140a^+^ osteolineage progenitor cells. (A) Gene commons analysis demonstrates that *Mx1*^+^*Ocn*^−^ P-SSCs (*MON*) and *Mx1*^+^*Nes*^+^ P-SSCs (*MNS*) highly express *Leptin receptor* (*Lepr*) and *Gremlin 1* (*Grem 1*) further demonstrating that these SSC populations share characteristics with previously studied BM-SSC populations. Further, KDR is found to be uniquely expressed in P-SSCs compared to other SSCs. (B) KDR^+^CD140a^+^ FACS analysis of P-SSCs (CD45^−^CD31^−^Ter119^−^ *Mx1*^+^*Ocn*^−^) and periosteal derived controls (CD45^−^CD31^−^Ter119^−^ *Mx1*^−^*Ocn*^+^) (C) KDR^+^CD140a^+^ FACS analysis of BM-SSCs (CD45^−^CD31^−^Ter119^−^*Mx1*^−^*Nes*^+^) and BM derived CD45^+^ controls. (D) Summary of FACS analysis demonstrates that *Mx1*^+^*Ocn*^−^ P-SSCs uniquely express KDR^+^CD140a^+^ (72%) compared to BM-SSCs and control populations (n = 3, p < 0.0001).

## Discussion

Herein, we sought to assess the functional genetic differences between mouse BM-SSCs and P-SSCs. These cell populations displayed differing apparent roles in fracture repair, so we hypothesized that their differences would be borne out in genetic expression analyses. We used *Mx1*^+^*Nes*^+^ cells from BM as BM-SSCs and *Mx1*^+^*Ocn*^−^ cells from periosteal tissues as P-SSCs. Using these cells, we performed a microarray analysis to compare their functional genetic differences. However, we were unable to find statistically significant difference in gene expression of these two populations. This is not unexpected considering that these both represent skeletal stem/progenitor cell populations, albeit from differing sources. On further analysis, we did find a novel marker for P-SSCs in comparison to BM-SSCs, which was KDR (aka VEGFR2, Fig. 4D). Additionally, there were other potential candidate genes upregulated in each SSC population in comparison to controls but their functional significance was unclear. Thus, while we did not find differential gene expression by clustered microarray analysis, we were able to find few unique genetic differences suggesting that these two cell populations may have subtle but critical functional differences.

Among the several markers previously demonstrated for SSCs, including Gremlin 1, Leptin Receptor and Nestin, we chose to isolate SSCs based on expression of *Mx1*. Unlike other markers, *Mx1*^+^ cells were shown to contribute to adult osteolineage cells by live *in vivo* imaging. This was further demonstrated here, which confirms their identity as osteolineage cells (Fig. 1A). Given *Mx1* marker has been known to label upstream hematopoietic lineage cells, we carefully isolated *Mx1*^+^ SSC populations by using SSC surface markers (CD105 and CD140a) and by negatively selecting against CD45^+^ hematopoietic lineage cells and CD31^+^ endothelial lineage cells as previously described. In addition we found *Mx1*^+^ cells are present in both the BM and periosteal tissue compartments, so we transiently label *Mx1*^+^ cells to isolate BM-SSCs and P-SSCs from each compartment, respectively. For BM-SSCs, *Mx1*^+^ cells were further purified by co-expression with *Nestin*^GFP^. By comparison, the P-SSC population was further purified by removing *Ocn*^GFP+^ adult osteolineage cells from the population. Inherently, our BM-SSC population was a more highly purified population than the P-SSC population used in this study, which is important to recognize for data interpretation. Still, both of these populations were found to express *Leptin Receptor* and *Gremlin 1,* showing that these populations are comparable to previously reported SSC populations, and this also supported our microarray findings.

We additionally isolated CD51^+^ cells as another population representing BMSCs for comparison to P-SSCs. This marker along with platelet derived growth factor-alpha had previously been shown to be expressed on *Nestin*^GFP+^ BM-SSCs. However, in our study, we found that this population was far different from the *Mx1*^+^*Nes*^GFP+^ BM-SSC population (Fig. 2G). In comparison, *Mx1*^+^*Nes*^+^ BM-SSCs and *Mx1*^+^*Ocn*^−^ P-SSCs were more closely related than CD51^+^ cells were than with either of these cell populations. This finding suggests that CD51^+^ cells may represent a distinct population cells than other BM-SSCs.

From our analysis, we identified KDR as a selectively expressed gene for P-SSCs compared to BM-SSCs. KDR is also known as VEGF receptor 2 (VEGFR2) and exerts its actions via binding VEGF. This receptor is known to be widely expressed on CD31^+^ endothelial cells. To minimize a potential endothelial contaminant in our cell isolation of P-SSCs, we had eliminated CD31^+^ cells during our collection making this less likely. Of note, it has been shown that human periosteal derived progenitor cells (PDPCs) display many characteristics of bone marrow MSCs and express VEGF receptor (Flt1 and KDR/Flk1) proteins [21]. Although the KDR expression in endogenous human PDPCs is not yet determined because they used *in vitro* cultured periosteal cells, our data showed that FACS-isolated murine P-SSCs have selective expression of KDR on their surface, supporting the possibility of KDR as a selective marker for P-SSCs and the relevance of our gene expression analysis. As a verification step, we performed pooled microarray analysis using gene commons data and also performed FACS analysis on our cells (Fig. 4). Both of these analyses confirmed significant upregulation of KDR on *Mx1*^+^*Ocn*^−^ P-SSCs compared to BM-SSCs (Fig. 4B & D). With this in mind, the expression of KDR on *Mx1*^+^*Ocn*^−^ P-SSCs is interesting because P-SSCs are believed to rapidly react to bone injuries and it would represent an efficient control mechanism for both endothelial cells and P-SSCs to respond to the same signaling molecule. Thus, during states of injury or inflammation, both cells would become activated in part for an angiogenic process and in part to initiate bone repair process, which inherently go hand-in-hand. Of further note is that periosteal tissue is known to be highly vascularized, and angiogenesis likely proceeds from the periosteal tissue. In either case, we would hope to further explore KDR as a potential regulatory mechanism of P-SSCs.

In summary, we performed a microarray analysis on mouse *Mx1*^+^*Nes*^+^ BM-SSCs and *Mx1*^+^*Ocn*^−^ P-SSCs and found that these are a similar population of cells without apparent differences readily assessed by gene expression analysis. However, our scatter plot analysis did show potential differences in gene expression although it did not reach statistical significance. The inability to find differential gene expression may be related to the residual heterogeneity of the cell populations. Still, both populations were found to express *Leptin Receptor* and *Gremlin 1*, which is consistent with their findings as SSCs and also supported the microarray analysis. We also found an interesting uniquely expressed gene in P-SSC, which was KDR. While the significance of this is yet to be determined, it represents an interesting gene because of its relationship to endothelial cells and the angiogenic response and the fact that periosteum is a highly vascularized tissue. Other studies to explore would be single cell analysis or exploring the possibility of environmental cues as the basis for the different functional roles between BM-SSCs and P-SSCs.

## Acknowledgments

Research reported in this publication was supported by National Institute of Arthritis and Muscloskeletal and Skin Diseases of the National Institutes of Health under grant number 1K01AR061434 and by the Bone Disease Program of Texas Award to D.P and The Caroline Wiess Law Fund Award to D.P.

## Supporting Information

**Supplemental Table 1.**
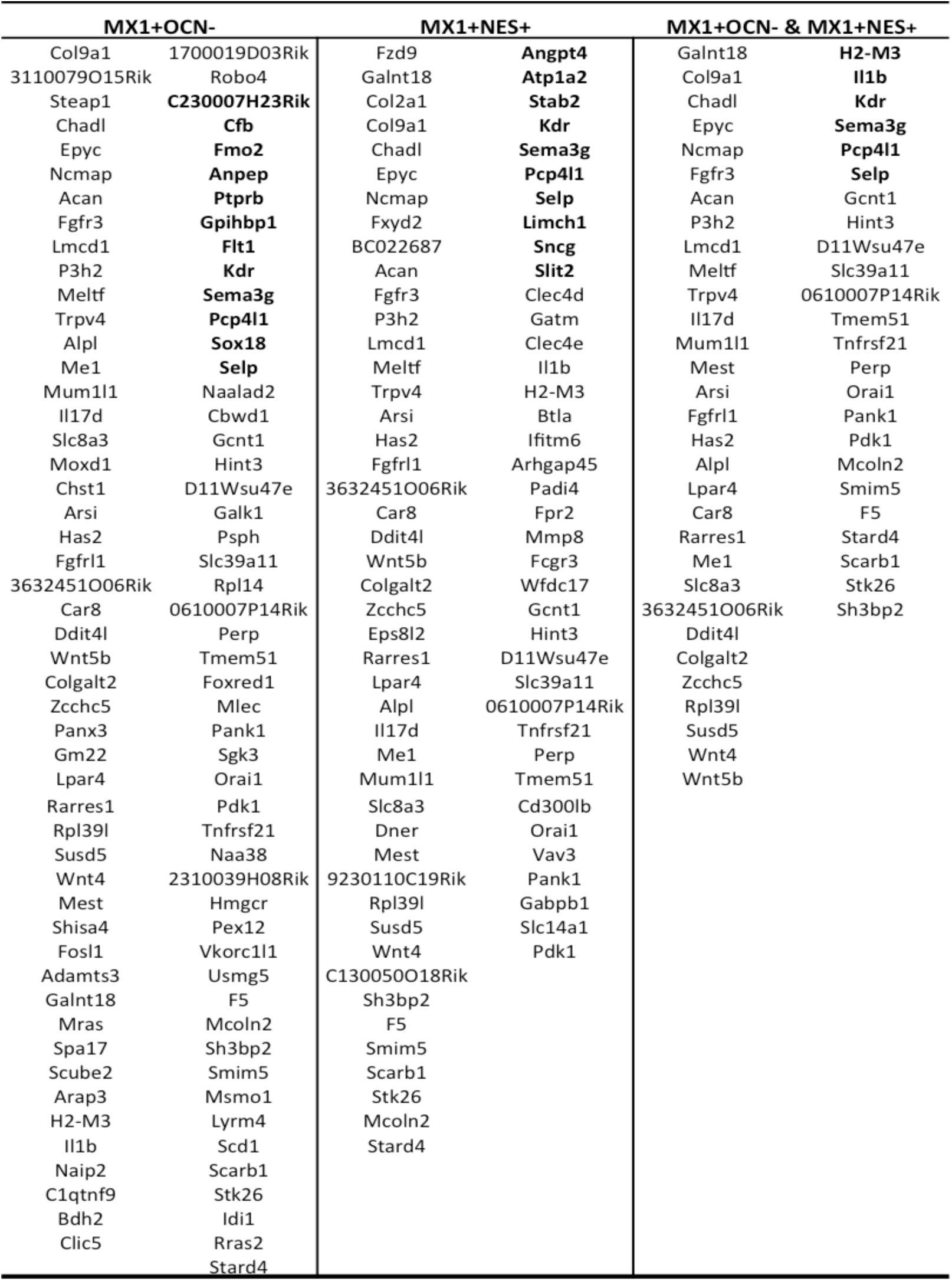
All differentially expressed genes comparing P-SSCs and BM-SSCs with both *CD45*^+^ cells and *OSX*^+^ cells (p < 0.05).

**Supplemental Table 2.**
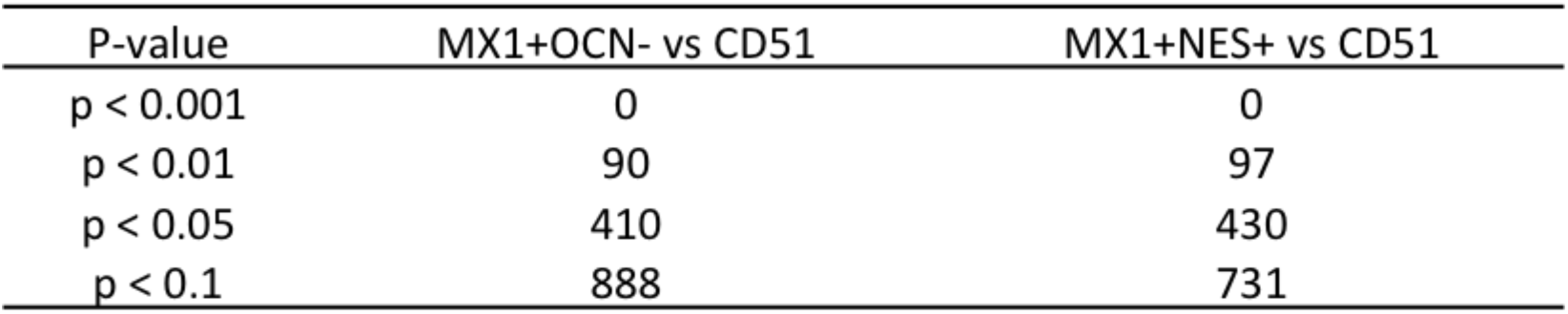
Table showing number significantly different genes and p-values.

| P-value | MX1+OCN- vs CD51 | MX1+NES+ vs CD51 |
| --- | --- | --- |
| p < 0.001 | 0 | 0 |
| p < 0.01 | 90 | 97 |
| p < 0.05 | 410 | 430 |
| p < 0.1 | 888 | 731 |

